# Copper resistance predicts heritable transgenerational fitness variation in the clonal duckweed *Spirodela polyrhiza*

**DOI:** 10.1101/2025.07.03.662999

**Authors:** Alexandra Chávez, Martin Schäfer, Iris Finkemeier, Shuqing Xu, Meret Huber

**Author notes:** Correspondence: Meret Huber, Johann-Joachim-Becher-Weg 7, 55128 Mainz, Germany.

## Abstract

Transgenerational plasticity in the absence of genetic change can alter organismal fitness, yet we know little about the genetic basis of transgenerational fitness effects. Here, we explored whether transgenerational fitness effects can be predicted by the rate of vegetative reproduction, plant resistance and defence. Thereto, we exposed monoclonal, single-descendant lineages from 56 globally distributed genotypes of the clonal duckweed *Spirodela polyrhiza* to five generations of copper excess, followed by five generations without stress, and then measured plant fitness and phenotypes in both environments. Previous copper excess reduced fitness variation within each genotype and elicited heritable and reproducible fitness effects that depended on both the genotype and environment: genotypes that benefited from previous stress under recurring conditions suffered when the stress was absent, and vice versa. These transgenerational fitness effects were predicted best by copper resistance, explaining 30% of the intraspecific variation: the more susceptible a genotype, the more its offspring suffered from ancestral stress in the absence of recurrent stress; however, these offspring became more resistant to copper excess, likely by protecting the photosystem II. Together, these findings show that transgenerational fitness effects are heritable and variable and likely controlled by genes that regulate plant reproduction and defence.

## INTRODUCTION

Global change accelerates environmental fluctuations. Thus, the ability to retain stress memory and pass environmental cues or traits to offspring may boost species resilience. Indeed, phenotypes can change across generations in the absence of genetic alterations [1]. Most of these changes appear only in immediate offspring –these changes are therefore referred to as maternal effects [1, 2]. More rarely, environmental cues trigger variations that persist for two or more generations, the latter referred to as transgenerational plasticity [3-8]. Notably, transgenerational plasticity can vary among genotypes, suggesting a genetic basis [3, 4, 9].

Elucidating the genetic basis of transgenerational plasticity has proven to be challenging. A recent study showed that across *Arabidopsis thaliana* genotypes, transgenerational effects in gene expression correlates with the abundance of several transposon classes [3]. However, such correlations remain rare, likely because of two hurdles: First, one must confirm that observed phenotypes stem from transgenerational plasticity rather than genetic change. This task requires following single descendants of clonal or highly inbred lineages over several generations [10]. Second, one must quantify phenotypes and organismal fitness in multigenerational experiments using dozens of genotypes. Such large-scale and long-term experiments are not feasible in most higher plants.

The first step in identifying the genetic basis of transgenerational plasticity is to quantify the proportion of variance in these transgenerational effects attributed to genetic differences. Although an increasing number of studies shows that species can be transgenerationally plastic [3-8], heritability estimates of transgenerational plasticity are scarce [11]. Heritability of a trait can be estimated as broad-sense heritability, which explains which fraction of the phenotypic variance is explained by all genetic effects [12]. Such broad-sense heritability are relevant for species reproducing asexually [13]. Heritability can also be estimated as narrow-sense heritability, in which only the additive genetic effects are considered [12]. Narrow-sense heritability is especially important in sexual reproduction because it represents the proportion of phenotypic variance attributable to additive genetic effects, the components most reliably transmitted from parents to offspring [13]. Estimating heritability, either broad-or narrow-sense, is key to infer whether the genetic basis of a trait can be identified.

Although often assumed to be adaptive, transgenerational plasticity is not necessarily beneficial. For example, if stress-induced transgenerational plasticity reduces offspring provisioning, then offspring fitness would decrease regardless of the environment, a phenomenon known as the silver-spoon effect [14]. Conversely, if stress-induced transgenerational plasticity triggers defences that are transmitted to offspring, these defences might incur costs under benign conditions due to the expense of maintaining vertically transmitted defences, but they could be advantageous under recurring stress. The phenomenon where mild stress benefits organisms, particularly when the stress recurs, is known as hormesis [15, 16]. Although hormesis is often studied within generations [17], its effects can persist across generations. For instance, *Caenorhabditis elegans* exposed to different abiotic stressors increased the lifespan and reproductive traits of non-exposed offspring even three generations after the ancestral stress when compared to the offspring of unstressed parents [9]. Additionally, hormesis may depend on the genetic background: for example, *Drosophila melanogaster* benefited from the application of dead fungal spores only in the absence of antifungal immunity [18]. These observations raise the questions whether transgenerational plasticity is ultimately beneficial or costly, and whether these outcomes may depend on the genotype and environment in which offspring develop.

Apart from altering a genotype’s average fitness, transgenerational plasticity may also affect the variability of plant fitness within a genotype. Often, transgenerational plasticity is assumed to increase phenotypic variation within a genotype, enabling certain offspring to succeed, a strategy known as diversified bet-hedging [19]. Diversified bed-hedging is supported from studies on maternal effects: for example, in several vertebrates and invertebrates, previous stress or fluctuating conditions increases egg size variation [20, 21]. However, it remains unclear whether transgenerational effects follow a diversified bet-hedging strategy or instead favour conservative bet-hedging [19], which reduces offspring fitness variation at the cost of maximal fitness.

One of the few flowering plants in which it is feasible to assess whether stress alters both the variance and mean of fitness across multiple generations–without genetic change–is the globally distributed aquatic giant duckweed, *Spirodela polyrhiza*. The plant consists of a flat, thallus-like shoot, the so-called frond, which float on the water surface. Under optimal conditions, the species reproduces very rapidly and exclusively vegetatively through budding every two days [22], thereby allowing precise, real-time measurements across many generations and genotypes. *S. polyrhiza* inhabits small, nutrient-rich water bodies and often endures fluctuating environmental stresses, such as copper excess [23]. Copper excess poses a major threat in aquatic ecosystems [24]; it primarily inhibits photosystem II, thereby triggering oxidative stress much like several herbicides, salt, and UV-light. To defend against oxidative stresses, *S. polyrhiza* produces glycosylated flavones, mainly apigenin and luteolin glucosides, as well as glycosylated and malonylated anthocyanins [4, 25, 26].

Using 56 world-wide sampled *S. polyrhiza* genotypes, we tested whether copper excess triggers transgenerational plasticity, thereby altering both the mean and variance in plant fitness across generations. Notably, we found that the magnitude and direction of these transgenerational fitness effects could be predicted by the copper resistance of each genotype, suggesting that genes controlling plant reproduction and defence govern transgenerational plasticity.

## MATERIALS AND METHODS

### Plant growth conditions

Plants were grown inside growth cabinets (GroBank, CLF PlantClimatics, Wertingen, Germany) at 150 μmol photons m^-2^ s^-1^, 28°C and 16:8 hrs of light and dark cycle using sterilized N-medium, which supports optimal growth [27]. All experiments were conducted under axenic conditions. During single descendant propagation, we minimized biases of environmental variation within the growth cabinets by randomizing the rack order within each shelf and moving the racks from right to left every propagation day (every two to three days). To differentiate mother and offspring, we marked the mother fronds using a permanent marker (Stabilo OHPen Universal, Heroldsberg, Germany).

### Statistical analysis

All data was analysed in R version 4.4.0 [28]. To upload, organize and summarize data we used the packages readxl v1.4.3 [29], dplyr v1.1.4 [30], data.table v1.16.4 [31], tidyr v1.3.1 [32]. For mixed effects models, we used the package glmmTMB v1.1.10 [33] and DHARMa v0.4.7 [34] to verify the adjustment of the fitted models. We plotted all data with ggplot2 v3.5.1 [35].

### *Transgenerational experiment with 56* S. polyrhiza *genotypes*

#### Pre-treatment, recovery, fitness assay

To understand the genetic basis of transgenerational plasticity, we selected 56 worldwide-distributed *S. polyrhiza* genotypes, representing the four genetic clusters of this species (table S1). After surface sterilization with sodium hypochlorite and cefotaxime (methods S1), we acclimated the plants by growing them in 150 mL of N-medium within 250 mL Erlenmeyer flasks (Fisher Scientific, Schwerte, Germany). After one month of acclimation, we placed six fronds per genotype into separate 30 mL polypropylene tubes (Fisher Scientific, Waltham, USA) filled with 25 mL of N-medium and closed with sterilized foam plugs (CarlRoth, Karlsruhe, Germany) (figure S1a, table S2). Once the next generation fully developed, we initiated the “pre-treatment” by transferring the offspring into tubes containing 25 mL N-medium with or without copper excess (20 μM CuSO_4_) (N = 3 per genotype, figure 1a, figure S1b). Then, the plants were propagated as single descendants for five consecutive generations. Subsequently, the offspring of the fifth generation were moved to control conditions and propagated as single descendants for an additional five generations (“recovery”, figure 1a), reaching generation 10.

**Figure 1.**
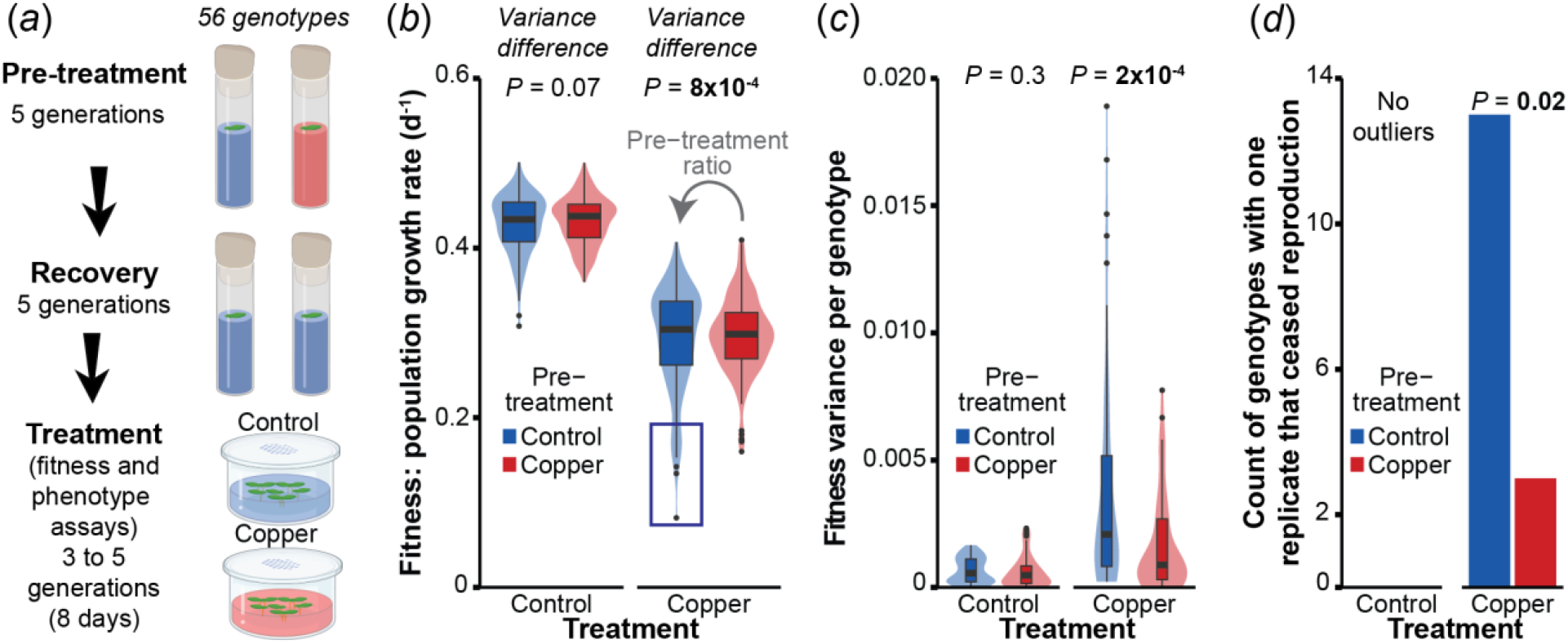
Ancestral copper excess in *Spirodela polyrhiza* reduces within-genotype variation in plant fitness under copper excess. (*a*) Experimental setup. Blue liquids refer to control condition and red liquids refer to copper excess. (*b*) Copper pre-treatment decreased overall variance in population growth rates under copper excess. Variance difference *P*-values, to compare homogeneity of variance between pre-treatments, refer to Levene tests for homogeneity of variance. N = 165-168. Each data point represents an individual sample. (*c*) Copper pre-treatment decreased the variance in population growth rates within genotypes. *P*-values to compare the variance between pre-treatments refer to Kruskal-Wallis rank sum tests. N = 56. Each data point represents one genotype. (*d*) Copper pre-treatment decreased the number of genotypes in which one replicate ceased reproduction (lower than 70% of the genotype’s median). No outliers refer to the absence of genotypes with one replicate ceasing reproduction. The *P*-value refers to a test of equal or given proportions.

Finally, we conducted fitness and phenotype assays: We placed the first offspring of the 10th generation in 250 mL transparent polypropylene beakers filled with 150 mL N-medium containing copper excess (20 μM CuSO_4_), and the second offspring in beakers filled with N-medium without copper excess (“treatment”). The beakers were covered with perforated transparent plastic lids (figure 1a, figure S1c). Plants were grown freely for eight days, corresponding to three to five generations, after which plants were harvested. All plant material, except the plants used to initiate the assay (“starting frond”), were briefly dried with a tissue paper, flash-frozen in liquid nitrogen, and stored at -80°C until metabolite extraction.

To assess plant fitness, we measured the increase in plant surface area across the eight days. Thereto, we photographed the plants at the beginning and end of the assays prior harvest using a camera box equipped with a webcam (HD Pro Webcam C270, Logitech, Lausanne, Switzerland; webcam software 2.12.8). Surface area was estimated using ImageJ 64 v5 [36].

To measure the maximal photosystem II quantum yield of dark-adapted plants, here defined as photosystem II efficiency (F_v_/F_m_ ratios), we collected the starting fronds after the eight-day fitness assay and placed each frond individually into 48-well plates (Sarstedt AG & Co. KG, Nümbrecht, Germany) filled with 1.2 mL of N-medium per well. Once the next generation had emerged under 110 μmol photons m^-2^ s^-1^, we removed the starting fronds, incubated the plates in darkness for 10 minutes within the climate cabinets and then transferred the plates to an IMAGING-PAM M-Series MAXI version (Heinz Walz, Amtsgericht Bamberg, Germany) to obtain the F_v_/F_m_ ratios (methods S2). The MAXI version included an IMAG-MAX/L measuring head, IMAG-K7 camera and mounting stand IMAG-MAX/GS.

#### Metabolite quantification

To assess the effect of copper pre-treatment on the accumulation of flavones and anthocyanins, we ground the plant material from all replicates to a fine powder using a MM301 Mixer Mill (Retch, Haan, Germany) and then pooled all samples to obtain mean values per genotype, pre-treatment and treatment (N = 56). 30 mg of the pooled plant material was extracted using acidified methanol. In these extracts, we subsequently quantified the flavones apigenin-7-O-glucoside, apigenin-8-C-glucoside, luteolin-7-O-glucoside and luteolin-8-C-glucoside by LC-MS, and the anthocyanins cyanidin-3-O-(6-O-malonyl-beta-glucoside) and cyanidin-3-O-glucoside–here referred as cyanidin-3-malony-glucoside and cyanidin-3-glucoside–by HPLC-PDA (methods S3). The peaks from all flavones, the LC-MS/MS data, were integrated with LabSolution Insight Version 4.0 SP6 (Shimadzu) and the peaks from anthocyanins, the HPLC-PDA data, with LabSolution Version 5.123 (Shimadzu, Duisburg, Germany).

#### Estimating fitness and pre-treatment ratios

We estimated daily population growth rates (“fitness”) by taking the natural logarithm of the ratio of the final plant surface area to the initial plant surface area and then dividing that value by the number of days the assay lasted (Formula S1) [37]. To assess whether genotypes differ in their transgenerational response, we calculated fitness pre-treatment ratios for each environment. These ratios represent the population growth rates of copper pre-treated plants relative to the mean population growth rates of control pre-treated plants in the respective environment (Formula S2) [5]. Finally, to assess genotype resistance, we calculated the ratio of population growth rates under copper excess to the genotype’s average population growth rates under control condition, using control pre-treated plants (Formula S3).

#### Effects of transgenerational plasticity on fitness variance

To assess whether copper pre-treatment alters the variance in population growth rates, we first determined whether the pre-treatment increased the overall variance and then examined if this change was due to increased variance within or across genotypes. To compare the homogeneity of variance between the two pre-treatments, we performed Levene tests with the package car v3.1.3 [38] for each environment separately. Similarly, to test whether the pre-treatment altered the variance among genotypes, we calculate mean values per genotype and performed Levene tests. To test whether the pre-treatment altered the variance within genotypes, we compared the variance between the two pre-treatments in each environment using Kruskal-Wallis rank sum tests implemented in R [28].

Furthermore, we counted the number of genotypes that contained one replicate with strongly reduced growth, defined as a reduction in population growth rates of at least 70%. We then assessed whether the pre-treatment altered the frequency of such genotypes using a test of equal or given proportions with the function prop.test of the package stats v4.4.0 implemented in R [28]. To determine whether copper pre-treatment affects the skewness of the population growth rate distributions relative to a normal distribution, we used the function test.skew from statpsych v1.7.0 [39] which employs Monte Carlo *p*-values. Finally, we compared distribution differences between pre-treatments using the Kolmogorov-Smirnov KS test with the function ks.test of the package stats v4.4.0.

#### Effects of transgenerational plasticity on fitness means

To assess the effect of the pre-treatment and treatment on the average of population growth rates, we used a linear mixed effect model defined as Fitness ∼Pre-treatment *Treatment +(1|Genotype). Similarly, to assess the effect of the treatment and genotype on the pre-treatment ratios, we used the model Fitness pre-treatment ratios ∼ Genotype *Treatment +(1|Rack), where “Rack” represents the physical structure of the replicates.

To estimate broad-sense heritability of the fitness pre-treatment ratios, we modified the function Cullis_H2 of the package remotes v0.1 (https://github.com/etnite/bwardr/) to incorporate glmmTMB models and validated the results using the package inti v0.6.6 [40]. To estimate narrow-sense heritability, we generated a kinship SNP matrix for 42 sequenced genotypes [41, 42] using Tassel v5 [43] and obtained heritability estimates with the package sommer v4.3.7 [44]. Finally, to assess whether transgenerational plasticity incurs environment-dependent fitness costs and benefits, we averaged pre-treatment ratios per genotype and environment and correlated these mean pre-treatment ratios across the two environments using the model Fitness pre-treatment ratios in control ∼ in copper +(1|Population).

#### Predicting fitness effects of transgenerational plasticity and inferring its physiological basis

To infer the genetic basis of the transgenerational plasticity, we first calculated the fitness and F_v_/F_m_ mean values per genotype considering naïve plants (plants under control pre-treatment), as well as the fitness and F_v_/F_m_ mean pre-treatment ratios per genotype (“genotype-averaged”). The genotype-averaged values of flavonoids were obtained when pooling the samples before the extractions. Second, we correlated the genotype-averaged fitness pre-treatment ratios for each environment separately with the genotype-averaged population growth rates under control, plant resistance, and the accumulation of flavones and anthocyanins of naïve plants, using linear models. To determine whether genotypes that benefit from ancestral stress under recurring conditions protect their photosystem II, we correlated the genotype-averaged fitness pre-treatment ratios with the genotype-averaged pre-treatment ratios of photosystem II efficiency (F_v_/F_m_) using the mixed effect model Pre-treatment ratio of fitness ∼ Pre-treatment ratio of F_v_/F_m_ * Treatment +(1|Genotype). In addition, we correlated the genotype-averaged pre-treatment ratios of fitness and F_v_/F_m_ for each environment separately using linear models.

To infer whether flavonoids protect photosystem II and thereby benefit plant fitness, we correlated genotype-averaged F_v_/F_m_ ratios and genotype-averaged population growth rates with the concentrations of the major *S. polyrhiza* flavones and anthocyanins, all in naïve plants, using the mixed effect model F_v_/F_m_ ∼ Metabolite concentration * Treatment + (1|Plate), as well as Population growth rates ∼ Metabolite concentration * Treatment +(1|Plate), where “Plate” denotes the batch of samples from which metabolites were extracted. Additionally, to assess the effect of the pre-treatment and treatment on anthocyanin concentrations, we used the mixed effect model Anthocyanin concentration ∼ Pre-treatment *Treatment +(1|Genotype).

#### Repetition experiment with 16 S. polyrhiza genotypes

To assess the reproducibility of the transgenerational fitness effects, we repeated the above-described experiment using 16 genotypes that spanned the entire range of fitness pre-treatment ratios (table S1). First, we confirmed that the genotypes were sterile at the beginning of the experiment by sugar tests (methods S1). Next, we pre-cultivated the plants in 250 mL Erlenmeyer flasks (Fisher Scientific) filled with 150 mL N-medium. The plants were then acclimated to the conditions of the tubes by placing eight single fronds per genotype into 30 mL polypropylene tubes (table S3) and propagating them as single descendants for three generations.

To initiate the pre-treatment, we dived the first and second offspring equally between copper excess (20 μM CuSO_4_) or control conditions (pre-treatment, figure 1a, table S4). We then propagated the plants for five generations in the respective pre-treatment environment, followed by five generations under control conditions (recovery, figure 1a). Finally, we transferred the first and second offspring (table S4) into 250 mL transparent polypropylene beakers filled with 150 mL N-medium to conduct eight-day fitness and phenotype assays, as described above. Thereto, half of the first and second offspring were placed in beakers containing copper excess (20 μM CuSO_4_), while the other half were placed in beakers under control conditions (“treatment”, N = 8, figure 1a).

To measure plant fitness and pre-treatment ratios, we followed the same procedure as described for the 56 genotypes. To assess whether the genotype-specific transgenerational fitness effects are reproducible, we correlated the standardized beta coefficients of the 16 genotypes from the first and second experiment. We extracted the regression coefficients from the model pre-treatment ratios of fitness ∼Genotype *Treatment +(1|Replicate per genotype)

+(1|Rack) for each experiment and then standardized the regression coefficients by multiplying by the sample standard deviation of X and dividing by the sample standard deviation of Y, making the coefficients comparable across predictors. To extract the standardized beta coefficients, we modified the stdCoef.merMod function (https://github.com/jebyrnes/ext-meta) to support glmmTMB models.

## RESULTS

### Previous copper excess alters both the mean and the variance in plant fitness

To test whether ancestral copper excess alters plant fitness, we grew single-descendant lineages of 56 world-wide distributed *S. polyrhiza* genotypes (table S1) for five generations under either control conditions or copper excess (pre-treatment). After five successive generations without stress, we measured population growth rate–using the increase in surface area as a proxy for plant fitness–and assessed plant morphology under both control conditions and recurrent copper excess. (figure 1a). First, we tested whether ancestral copper excess alters the variability of population growth rates. When considering all replicates, control pre-treated plants displayed a greater variance in population growth rates than copper pre-treated plants, particularly under copper excess (control conditions: *P* = 0.07; copper excess: *P* = 8×10^−4^, Levene test, figure 1b). This difference in variance among the pre-treatments arose from increased variability within genotypes rather than among them: under copper excess but not control conditions, within-genotype variation was higher in control than copper pre-treated plants (*P* = 2×10^−4^, Kruskal-Wallis rank sum test, figure 1c). In contrast, in either environment, the variance in mean population growth rates per genotype, i.e. the variance between genotypes, did not differ among the pre-treatments (*P* > 0.2, Levene tests, figure S2a-b).

The within genotype-variance was higher in control than copper pre-treated plants because several control pre-treated plants ceased reproduction under copper excess: when pre-treated with control condition, 12 genotypes included a replicate in which population growth rates was reduced by at least 70% compared to the genotype median, whereas when pre-treated with copper excess, only three genotypes showed such a reduction (*P =* 0.02, test of equal or given proportions, figure 1d, figure S2c). Consequently, under copper excess, the growth rate distributions were left-skewed in control but not copper pre-treated plants (copper: *P* = 0.03, Kolmogorov-Smirnov test, figure S2d). Under control conditions, population growth rates of all replicates remained similar. These patterns held even when we adjusted the threshold for reduced growth from 60% to 75% (table S5). Taken together, these data show that ancestral copper exposure narrows fitness differences within genotype under stress by preventing some replicates from ceasing reproduction.

Second, we tested whether ancestral copper excess alters the average fitness of genotypes. For each genotype and environment, we computed fitness pre-treatment ratios, which are the population growth rates of copper pre-treated plants relative to the mean population growth rates of control pre-treated plants. Fitness pre-treatment ratios depended on the genotype and environment: under control conditions, copper pre-treatment caused only marginal changes in fitness, ranging from a reduction of 10% to an increase of 15%, with 90% of the genotypes showing changes within ±8%. Under copper excess, the effects of the copper pre-treatment on fitness were larger, ranging from a reduction of 25% to an increase of 42%, with 90% of the genotypes varying within ±22% (figure 2a, figure S3). Consequently, depending on the genotype, transgenerational plasticity either increased or decreased plant fitness, with particularly strong effects under recurring stress.

**Figure 2.**
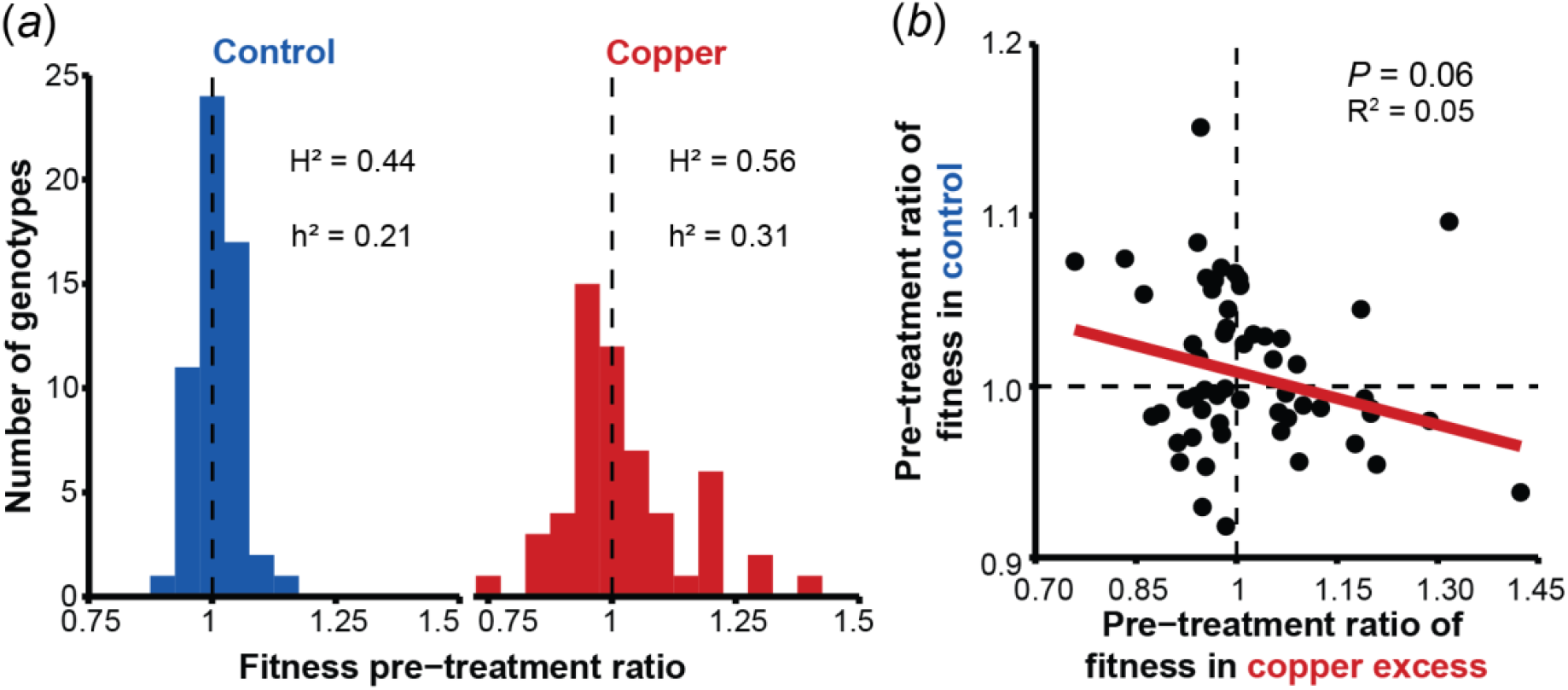
Copper-induced transgenerational fitness effects are heritable, variable across genotypes and environment-dependent, revealing a trade-off between enhanced performance under stress and reduced fitness under control conditions. (*a*) Histogram of the fitness pre-treatment ratios and their estimates of broad-sense (H^2^) and narrow-sense (h^2^) heritability. Fitness pre-treatment ratios represent the population growth rates of copper pre-treated plants relative to the mean population growth rates of control pre-treated plants, estimated over the 8-day fitness assay. N = 42-56. A pre-treatment ratio of 1, indicated by the dashed vertical line, signifies that ancestral stress had no effect on a genotype’;s mean fitness. (*b*) Scatterplot illustrating the trade-off in transgenerational fitness effects between environments: genotypes that benefited from copper pre-treatment under recurring stress exhibited reduced fitness when the stress was absent. *P*-value refers to a mixed effects model. Each data point represents the average fitness pre-treatment ratio per genotype, estimated over the 8-day fitness assay. N = 56.

To quantify the genetic proportion explaining variance in transgenerational fitness effects, we calculated both broad-(H^2^) and narrow-sense (h^2^) heritability on the fitness pre-treatment ratios. Under control conditions, heritability of the fitness pre-treatment ratios was moderate with H^2^ = 0.44 and h^2^ = 0.21. Under copper excess, both estimates increased, reaching H^2^ = 0.56 and h^2^ = 0.31. These data support the notion that genetic variation of transgenerational plasticity exists within species.

Genetic variation in transgenerational plasticity should evolve when it benefits plant fitness in one environment but reduces plant fitness in another environment. To test this hypothesis, we correlated each genotype’;s pre-treatment fitness ratio under control conditions with its corresponding ratio under copper excess. The pre-treatment fitness ratios tended to be negatively correlation between the two environments (*P =* 0.06, mixed effects model, figure 2b). In essence, genotypes in which ancestral stress improved their fitness under recurring stress suffered when the stress did not recur, and vice versa.

### Fitness effects of ancestral copper exposure can be predicted by copper resistance

To infer the genetic basis of transgenerational plasticity, we explored which factors predict within-species variation in transgenerational fitness effects by correlating the fitness pre-treatment ratios with population growth rates in control environments, plant resistance, and the production of defensive flavonoids. Plant resistance–defined as the relative reduction in population growth rates upon copper exposure–proved to be the best predictor of fitness pre-treatment ratios. In fact, variation in copper resistance explained roughly 30% of the variation in fitness pre-treatment ratios: the more susceptible a genotype was, the more its offspring suffered from ancestral stress under control conditions (*P =* 3×10^−6^, R^2^ = 0.28) but the better its offspring became protected from copper excess (*P =* 7×10^−8^, R^2^ = 0.33, figure 3a). Because susceptible genotypes reproduced faster in the absence of stress (*P =* 6×10^−6^, R^2^ = 0.27, figure 3b), fitness pre-treatment ratios could also be predicted by population growth rates, particularly under control conditions: the faster a genotype reproduced under control conditions, the more its offspring suffered from ancestral stress under those same conditions (*P =* 8×10^−8^, R^2^ = 0.53), yet its offspring were better protected under recurring stress (*P* = 0.05, R = 0.05, figure 3c). Individual defensive traits, namely the accumulation of antioxidative anthocyanins and luteolin glucosides, correlated only weakly positively with fitness pre-treatment ratios under control conditions and did not correlate to pre-treatment ratios under copper excess (figure S4). Overall, these findings reveal that intraspecific variation in transgenerational fitness effects can be predicted by differences in plant resistance and population growth rates.

**Figure 3.**
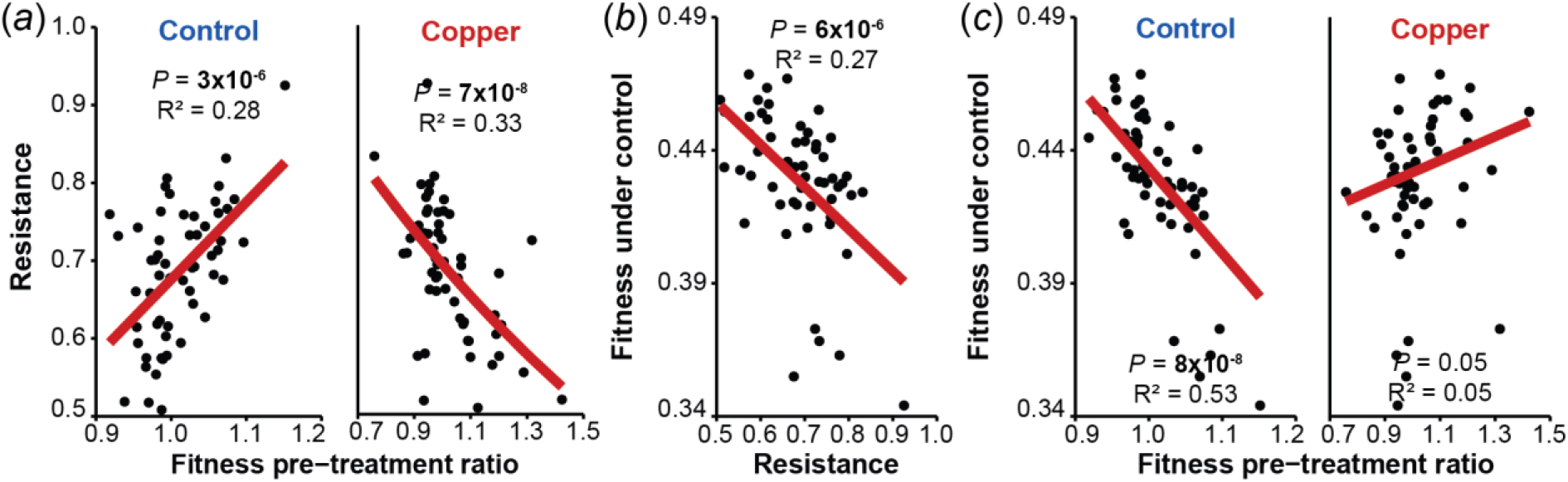
Within-species variation in transgenerational fitness effects were predicted best by copper resistance. (*a*) The fitness pre-treatment ratios (population growth rates of copper pre-treated plants relative to the mean population growth rates of control pre-treated plants) correlated positively with copper resistance under control conditions and negatively under copper excess. Copper resistance is defined as the relative reduction in population growth rates of control pre-treated plants under copper excess. (*b*) Genotypes with high population growth rates (“fitness”) under control conditions were susceptible to copper excess. (*c*) Genotypes with high reproductive rates (“fitness”) under control conditions suffered from ancestral stress if the stress did not recur; however, their offspring became more resistant to copper excess. *P*-values in a-c refer to linear models. Each data point represents a genotype’s mean values, estimated over the 8-day fitness assay. N = 56.

### Maintenance of photosystem II efficiency likely mediates transgenerational fitness effects

As copper excess leads to oxidative stress in plants, we examined whether genotypes that benefit from ancestral stress under recurring conditions also protect their photosystem II under stress. Thereto, we correlated the pre-treatment ratios of plant fitness with the pre-treatment ratios of F_v_/F_m_ values, a proxy for photosystem II efficiency. Under copper excess, the pre-treatment ratios of plant fitness correlated positively with those of F_v_/F_m_, whereas under control conditions, the pre-treatment ratios did not correlate (Control: *P =* 0.7; Copper: *P =* 0.02; *P*(F_v_/F_m x_Treat) = 9×10^−5^, mixed effects models, figure 4a). In other words, offspring benefiting from ancestral stress under recurring conditions also protected their photosystem II from oxidative damage.

**Figure 4.**
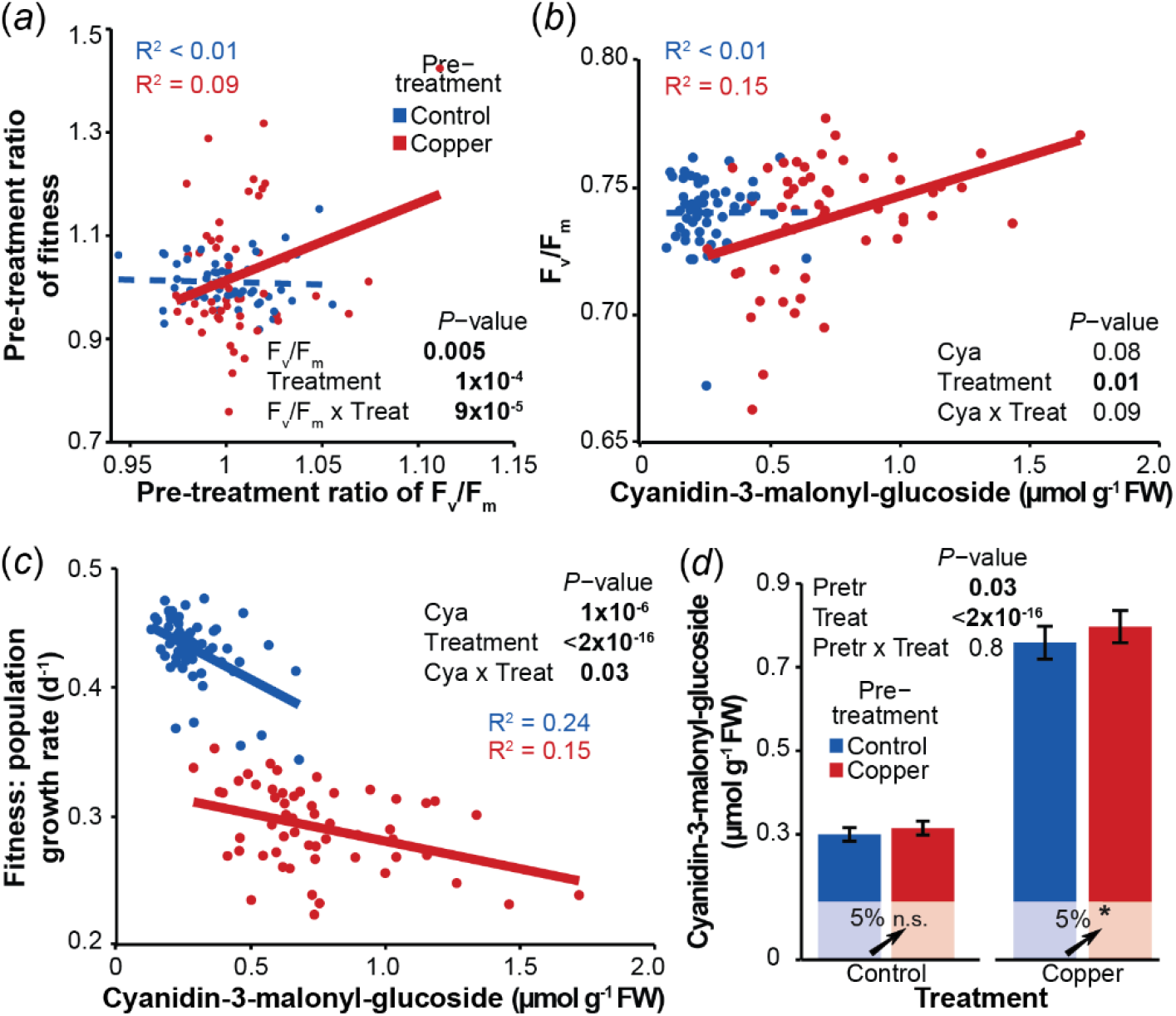
Genotypes that benefit from copper pre-treatment under recurring stress protect their photosystem II from oxidative damage, potentially through transgenerationally inherited anthocyanins. (*a*) The pre-treatment ratios of the photosystem II efficiency (pre-treatment ratio of F_v_/F_m_) correlate positively with the pre-treatment ratio of fitness under copper excess but not under control conditions. Pre-treatment ratios refer to the ratio in F_v_/F_m_ or plant fitness of copper pre-treated plants relative to the mean values of control pre-treated plants. Plant fitness is defined as daily population growth rates. (*b*) Genotypes with higher concentrations of cyanidin-3-malonyl-glucoside had higher photosystem II efficiency (F_v_/F_m_) under copper excess but not under control conditions. (*c*) Cyanidin-3-malonyl-glucoside levels correlated less negatively with population growth rates under copper excess than under control conditions. For (*a*)-(*c*): Each data point represents a genotype’s mean value. *P*-values refer to mixed effects models. Solid lines indicate correlations with *P* < 0.05, whereas dotted lines indicate correlations with *P* > 0.05. Population growth rates and metabolite levels were measured after eight days of fitness assay. F_v_/F_m_ values were measured from an offspring that emerged immediately after the 8-day fitness assay from the plant used to initiate the fitness assay. (*d*) Copper pre-treatment transgenerationally enhanced cyanidin-3-malonyl-glucoside concentrations regardless of the subsequent treatment. Error bars denote standard errors. *P*-values on top of the panel refer to a mixed effect model and *P*-value results within bars refer to mixed effects models, with n.s. for *P* > 0.05, and * for *P* < 0.05. Treat = treatment, Cya = cyanidin-3-malonyl-glucoside, Pretr = pre-treatment. N = 56.

**Figure 5.**
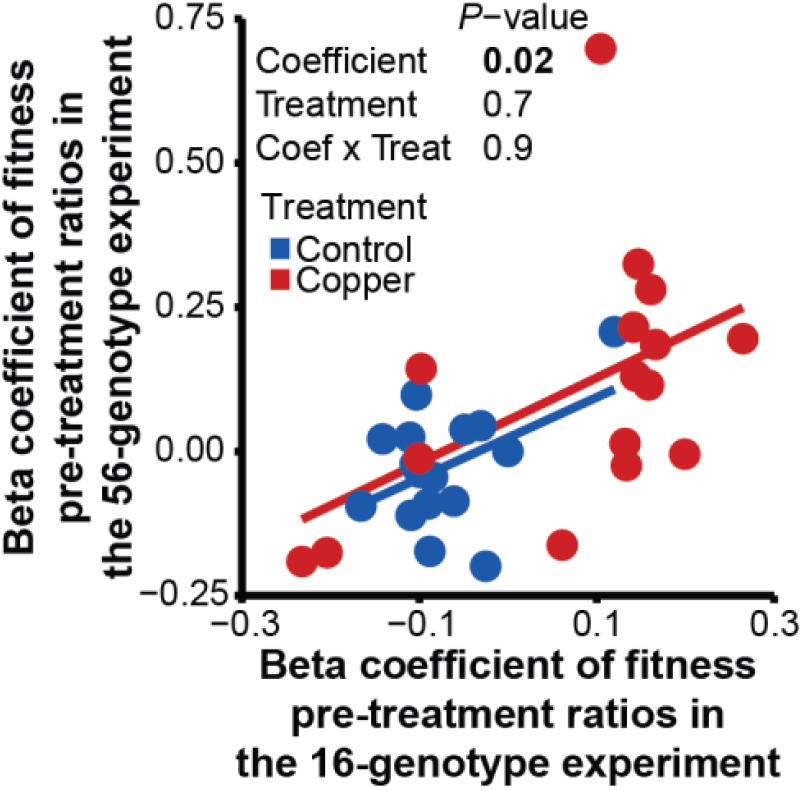
Genotype-specific transgenerational fitness effects are reproducible. Standardized beta coefficients, which quantify the effect of each genotype on the fitness pre-treatment ratio, correlated positively across two independent experiments. Each data point represents a genotype. *P*-value refers to a regression model. Coefficient = beta coefficient of fitness pre-treatment ratios, Coef = coefficient, Treat = treatment. N = 16.

We then assessed whether anthocyanins and flavones protect photosystem II and benefit plant fitness under copper excess. Among the 56 genotypes, the level of anthocyanins, but not flavones, correlated positively with F_v_/F_m_ ratios under copper excess but not control conditions (Cyanidin-3-malonyl-glucoside: Control: *P =* 0.9, Copper: *P =* 0.002; Cyanidin-3-glucoside: Control: *P =* 0.9, Copper: *P =* 0.01; mixed effects models, figure 4b, figure S5a). Moreover, the levels of these anthocyanins correlated strongly negatively with population growth rates under control conditions, and only weakly negatively under copper excess, indicating that anthocyanin production is costly under control conditions but beneficial under copper excess (*P*(Anthocyanins _x_Treat) < 0.03, mixed effects models, figure 4c, figure S5b). Collectively, these data suggests that the anthocyanins protect photosystem II and thereby benefit plant fitness under copper excess.

Next, we tested whether the genotypes that benefit from previous stress under recurring conditions retain elevated, stress-induced levels of anthocyanins across generations. On average, first-time copper excess doubled the levels of cyanidin-3-malonyl-glucoside and cyanidin-3-glucoside (*P* < 8×10^−10^, mixed effects model, figure 4d, figure S7a). Moreover, copper pre-treated plants retained 5% higher anthocyanin levels under both control conditions and copper excess (Cyanidin-3-malonyl-glucoside: *P*(Pre-treatment) *=* 0.03; Cyanidin-3-glucoside: *P*(Pre-treatment) *=* 0.06; mixed effects models, figure 4d, figure S7b). However, the pre-treatment ratios of both cyanidins did not correlate with the pre-treatment ratios of F_v_/F_m_ values or those of plant fitness (figure S7c-d). These findings suggest that either our metabolite analysis occurred at an unsuitable time or that other physiological mechanisms underlie the protection of photosystem II under recurring conditions.

### Transgenerational fitness effects are reproducible

To test whether our transgenerational effects in plant fitness are reproducible, we repeated the transgenerational experiment (figure 1a) with a subset of 16 genotypes that spanned the entire range of pre-treatment ratios observed under control conditions and copper excess (figure S8). To assess the reproducibility of our experiments, we compare the effect sizes of the explanatory variables across the two experiments. Thereto, we calculated the standardized beta coefficients for the pre-treatment ratios of each genotype. The beta coefficients from the two experiments correlated positively under both control and copper excess (*P*(beta coefficients) = 0.02, linear model, figure S6), showing that genotype-specific pre-treatment ratios of plant fitness are reproducible.

## DISCUSSION

Transgenerational plasticity can influence organismal fitness in a genotype-dependent manner, yet its genetic basis remains largely unknown. In our study, copper excess produced genotype-and environment-specific effects on offspring fitness after five clonal generations without stress. Notably, this intraspecific variation could be predicted by population growth rates and plant resistance, suggesting that genes governing plant reproduction and defence underlie transgenerational plasticity in *S. polyrhiza*.

Transgenerational plasticity varies among genotypes in several species, including *S. polyrhiza, Trifolium repens*, and *A. thaliana* [3, 4, 6], suggesting a genetic basis. Yet, identifying the genetic basis has proven to be challenging. One notable example is the work of Lin X, Yin J, Wang Y, Yao J, Li QQ, Latzel Vet al. [3], which showed that across *A. thaliana* genotypes, transgenerational variation in gene expression correlates with the abundance of several transposon classes. Here, by screening 56 *S. polyrhiza* genotypes, we found that transgenerational fitness effects can be predicted by copper resistance, explaining 30% of the intraspecific variation: more susceptible genotypes produced offspring that suffered from the ancestral stress under control conditions; however, these offspring became more resistant to copper excess. These strong genetic correlations suggests that genes involved in reproduction and defence underlie the fitness effects of transgenerational plasticity.

Two non-exclusive mechanisms could explain why susceptible genotypes produced offspring that suffered from ancestral stress under control conditions but became better protected to recurring copper excess. First, under stress, resistant genotypes might supply more resources to their offspring than susceptible genotypes, boosting offspring performance regardless the environment, a phenomenon known as the silver-spoon effect [14]. However, if this mechanism dominated, we would expect the offspring of resistant genotypes to be better prepared for recurring stress as well, which was not the case. Instead, our data support a second mechanism, referred to as hormesis [15, 16]: in susceptible genotypes, stress may trigger a stronger induction of stress signals or defensive substances than in the resistant genotypes. If the concentrations of these molecules are transgenerationally elevated, the offspring may perform worse under control conditions, due to the cost of defence [45], but better under recurring stress. To distinguish between these hypotheses, the silver-spoon effect and hormesis, one could increase the stress level in resistant genotypes and assess whether their offspring show enhanced protection against recurring stress.

To identify traits that are transgenerationally plastic and protect susceptible genotypes under recurring stress, we screened candidate defences and photosystem II efficiency. Genotypes that benefited from previous stress under recurring conditions also maintained superior photosystem II efficiency under copper excess, suggesting that photosystem II protection is a key factor mediating transgenerational fitness effects. However, the underlying physiological mechanisms remain largely unclear. We hypothesized that anthocyanins might play a role. Indeed, our data indicate that anthocyanins protect plant fitness and photosystem II efficiency under copper excess, consistent with their well-known role as antioxidants [46-49] and protectors of the photosystem II [50-52]. Furthermore, copper-induced anthocyanin levels were passed on to the next generation. However, intraspecific variation in the transgenerational inheritance of anthocyanins did not correlate with variation in photosystem II efficiency or plant fitness. These patterns suggest that either our metabolite analysis was conducted at an unsuitable time point or other physiological mechanisms contribute to the protection of photosystem II under recurring stress. Future experiments should manipulate anthocyanin levels genetically to determine whether elevated levels protect photosystem II under recurring copper excess. Additionally, testing other recurring stresses such as herbicides, salt or UV-stress could reveal whether antioxidants and photosystem II protection contribute to transgenerational cross-resistance to oxidative stress, thereby clarifying the ecological relevance of transgenerational plasticity.

In our study, we found that previous stress altered not only the average plant fitness but also the variation in fitness within genotypes. Specifically, ancestral copper excess reduced the variance in population growth rates within genotypes under recurrent stress by preventing some replicates from ceasing reproduction, a phenomenon that aligns with conservative bet-hedging [19]. These results contrast with the proposed strategy of asexual and genetically depleted organisms to overcome environmental fluctuations by transgenerationally diversifying, rather than homogenizing, phenotypes and fitness [53-55], a process described as diversified bet-hedging [19]. However, modelling the evolution of maternal effects showed that reduced phenotypic variance in the offspring can potentially stabilize population sizes by minimizing extreme maladaptive phenotypes [56]. Because variation in fitness can influence long-term population growth rates [57] and rates of evolution [58, 59], our findings underscore the importance of assessing not only average fitness but also within-genotype variation when evaluating the evolutionary consequences of transgenerational plasticity.

Although our study focused on an aquatic species reproducing exclusively asexually during the experiments, we argue that the insights gained extend beyond this specific reproductive mode and plant system. First, asexual reproduction is common among flowering plants and lower animal phyla [60-65], and many invasive weeds, crops, and keystone species, including *S. polyrhiza*, reproduce asexually [66, 67]. Therefore, even if our findings primarily apply to mostly asexually reproducing organisms, the data show that transgenerational fitness changes can substantially affect the population dynamics of key stone species. Second, we uncovered a likely genetic basis for transgenerational plasticity by leveraging the experimental advantages of our system, particularly the rapid asexual reproduction. Only with this study system we were able to measure fitness across dozens of genotypes over multiple generations. These mechanistic insights are crucial for understanding the genetic bases of transgenerational plasticity and provide the starting point to test whether transgenerational fitness effects are governed by genes controlling reproduction and defence also in other species, including those that reproduce sexually.

Taken together, our data show that transgenerational plasticity affects both the average and the variability of plant fitness across generations. Importantly, we can now predict a genotype’;s transgenerational fitness effects from its resistance, suggesting that genes controlling plant reproduction and defence govern transgenerational fitness effects.

## ACKNOWLEDGEMENTS

We would like to thank Tom Lieth for assisting with the sterilization and harvesting of plants, Anne Schreyer for extracting and measuring the flavonoids, Annika Brünje for helping us set up the IMAGING-PAM software and for her valuable comments on the manuscript, and Marie Serwaty-Sárazová for standardizing the sterilization protocol.

## DATA AVAILABILITY STATEMENT

All raw data and R scripts for the analyses and plots within this study are deposited in https://github.com/Plant-Evolutionary-Ecology-Lab/Genetic-Base-Transgenerational-Plasticty.

