## Supplemental Material for "Copper resistance predicts heritable transgenerational fitness variation in the clonal duckweed *Spirodela polyrhiza*"

#### Figures

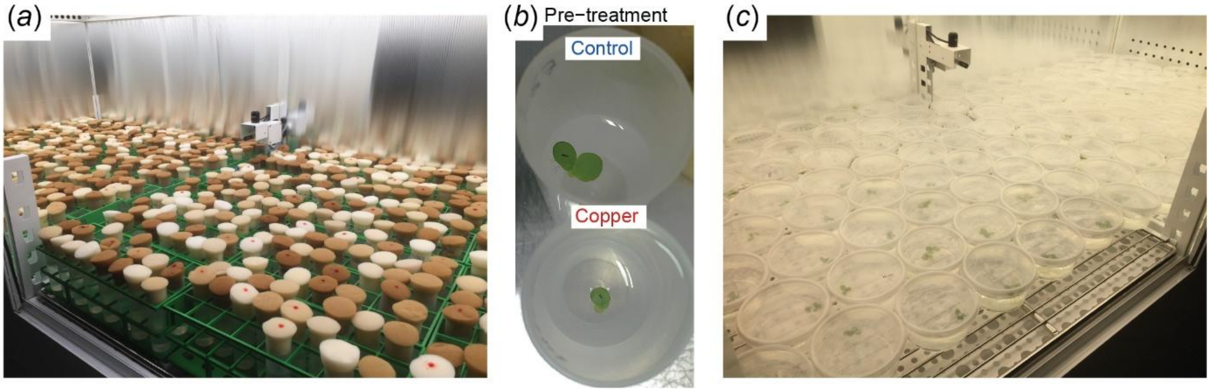


Figure S1. Distribution of samples within polypropylene tubes and beakers. (*a*) Plants within polypropylene tubes were distributed in racks, with maximum two generations per strain to homogenise the light incidence within the racks. Genotypes = 56, pre-treatments = 2, replicates = 3, generations = 2; 672 tubes. (*b*) Single descendant frond in control and copper excess during the pre-treatment phase. (*c*) Plants within polypropylene beakers during fitness and phenotype assays. Genotypes = 56, pre-treatments 2, treatments = 2, replicates = 3; 672 beakers.

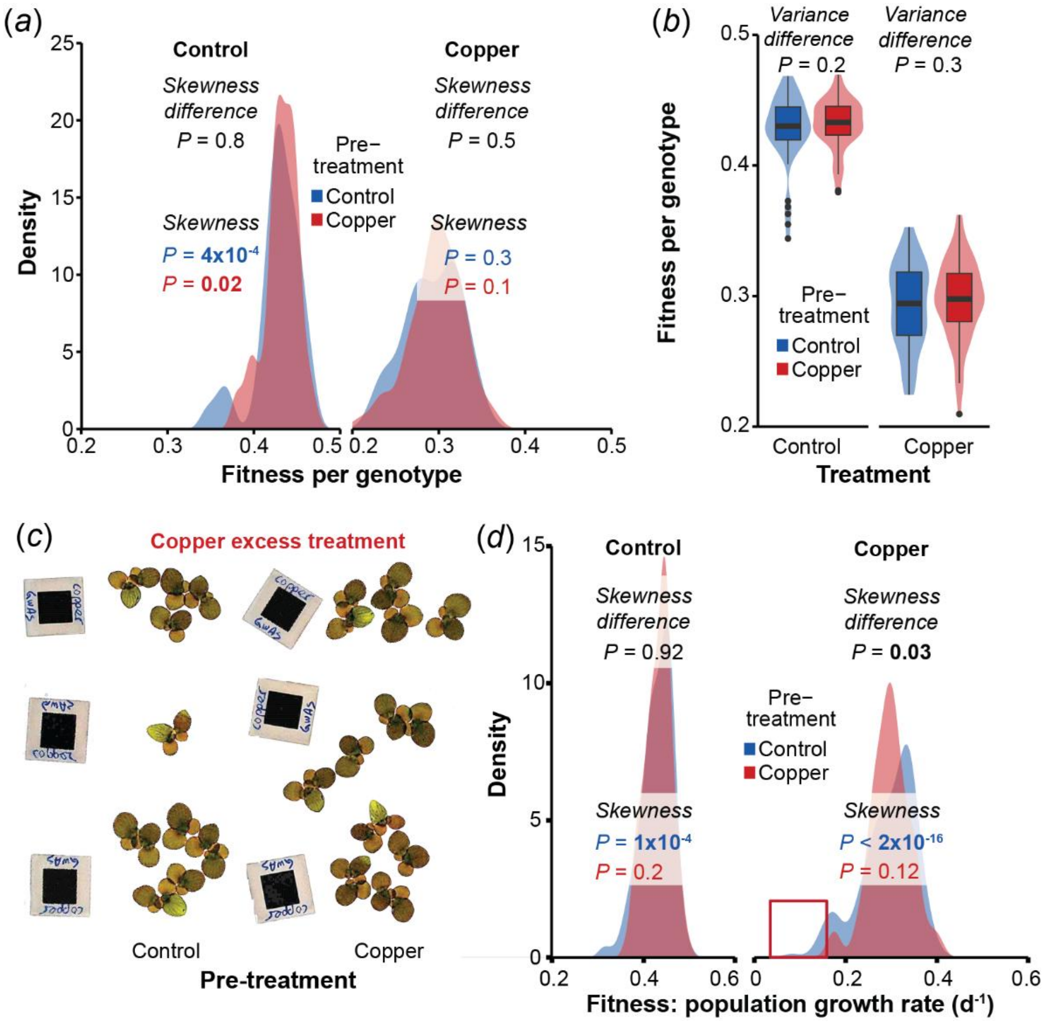


Figure S2. Under recurrent copper excess, ancestral stress exposure affects the distribution of samples within genotypes but not among genotypes. (*a*) Samples of genotypes exposed to copper excess for the first time showed stronger fitness decrease than samples from copper pre-treated genotypes. Skewness *P* values refer to the comparison of the data distribution to a normal distribution using Monte Carlo *p*-values. Skewness difference *P* values on top refer to Kolmogorov-Smirnov tests to compare the distribution between pre-treatments. Control and Copper headers refer to the treatment. Blue and red colour of the *P* values refer to control and copper pre-treatment, respectively. N = 165-168. (*b*) Variance between mean fitness per genotype of control and copper pre-treated plants remained similar under control and copper excess. Variance difference *P* values, to compare homogeneity of variances between pre-treatments, refer to Levene test for homogeneity of variance N = 56. (*c*) Genotype SP223, genotype with the strongest outlier under first time copper excess. (*d*) Mean fitness per genotype did not differ in their distributions when comparing control to copper pre-treated plants. Skewness *P* values refer to the comparison of the data distribution to a normal distribution using Monte Carlo *p*-values. Skewness difference *P* values on top refer to Kolmogorov-Smirnov tests to compare the distribution between pre-treatments. N = 56.

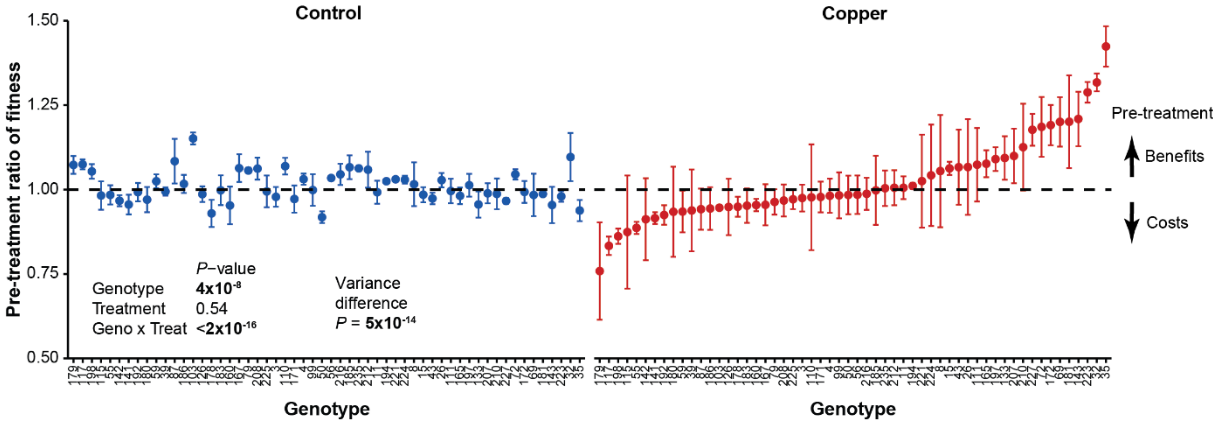


Figure S3. Pre-treatment ratios dependent on the treatment environment and genotype. *P* values on the left refer to a mix effects model; N = 3. Variance difference *P* value, to compare data homogeneity of variances between treatments, refers to Levene test for homogeneity of variance; N = 168. Control and Copper headers refer to the treatment.

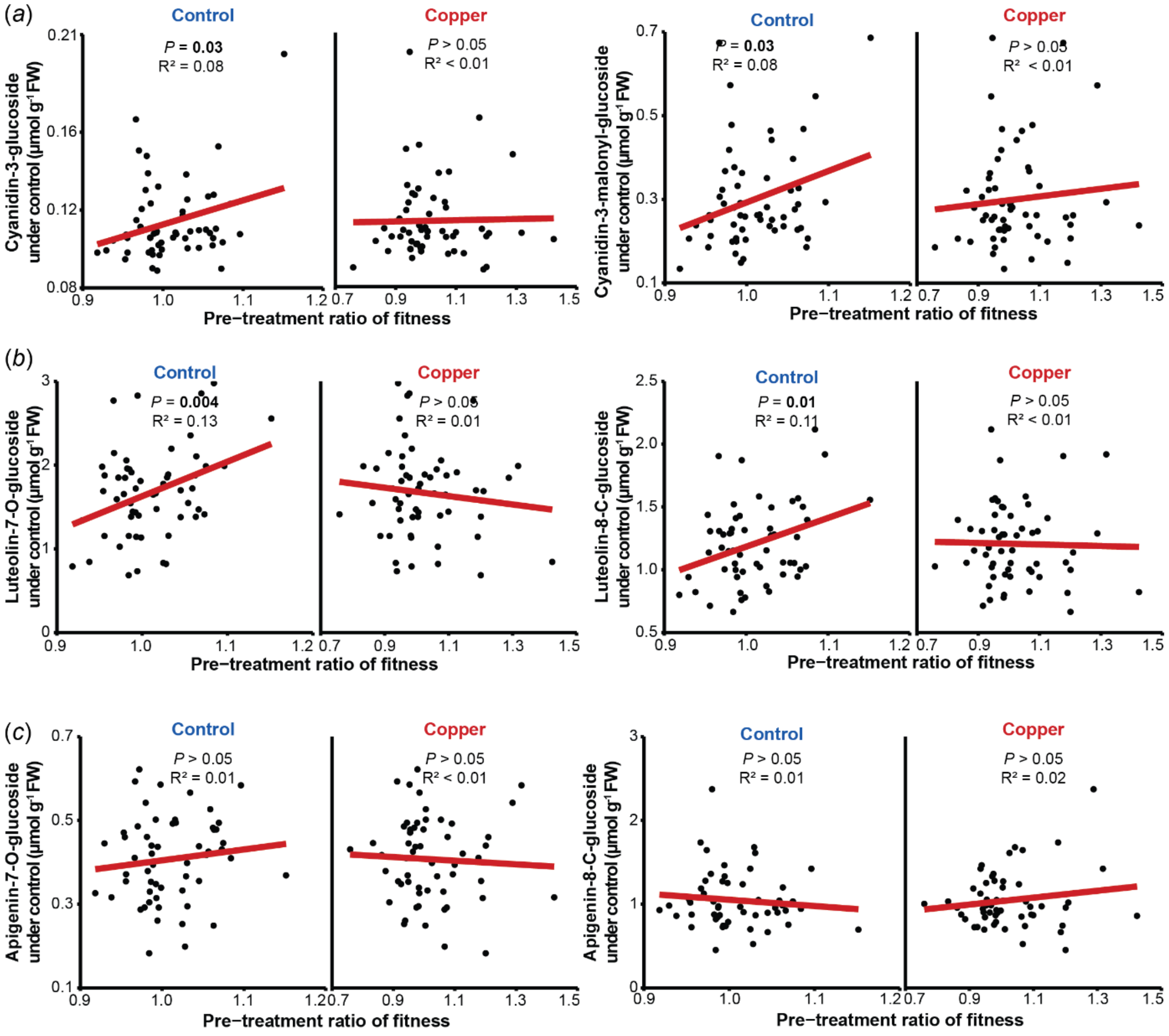


Figure S4. The offspring of genotypes with higher flavonoid concentrations do not have benefits or costs under recurrent copper excess. Genotype concentrations of (*a*) anthocyanins, (*b*) luteolins and (*c*) apigenins under control conditions did not predict the pre-treatment ratios of plant fitness (fitness of copper pre-treated plants relative to the mean fitness of control pre-treated plants). Mean flavone concentration per genotype was obtained through pooling of the genotype replicates before extraction of the metabolites. *P* values per treatment were calculated with simple linear regressions. Control and Copper headers refer to the treatment. N = 56.

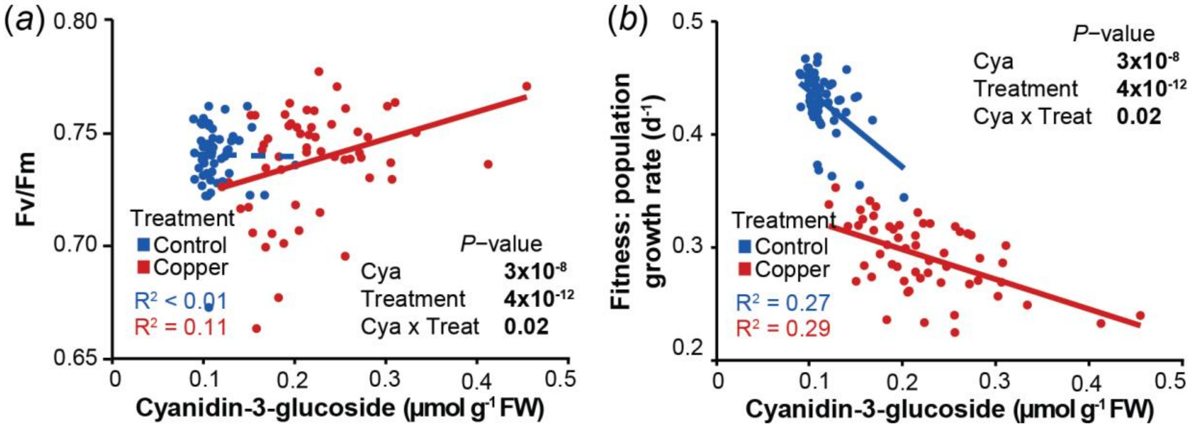


Figure S5. Genotypes with higher concentrations of cyanidin-3-glucoside show better photosystem II activity and less fitness costs. (*a*) Genotypes with higher concentrations of cyanidin-3-glucoside had higher photosystem II activity under copper excess, but no correlation was observed under control conditions. (*b*) The costs of cyanidin-3-glucoside in plant fitness were overcome under copper excess. Fv/Fm represents mean values per genotype. Mean cyanidin concentration per genotype was obtained through pooling of the genotype replicates before extraction of metabolites. Continuous lines refer to correlations with $P < 0.05$, and dotted lines refer to correlations with $P > 0.05$ from mixed effects models. *P* values refer to mixed effects models. *P* values refer to mixed effect models, where significant interactions ($Cyanidin_x$Treatment) show the correlation difference between treatments. N = 56.

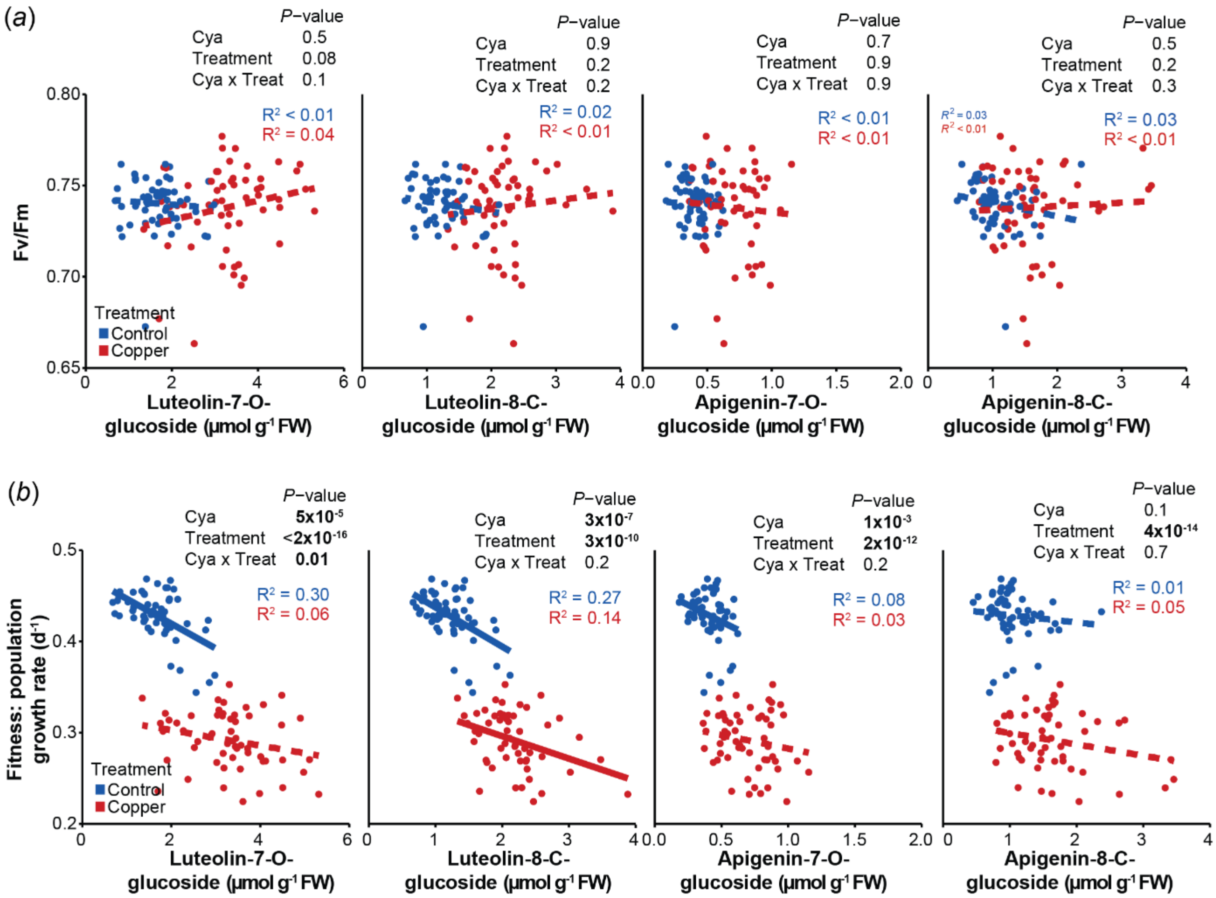


Figure S6. Accumulation of flavones did not represent benefits for plants under first time stress. (*a*) Photosystem II activity (Fv/Fm) did not correlate with flavone concentrations under copper nor control conditions. (*b*) Genotypes with higher flavone concentrations showed costs in plant fitness. Fv/Fm and fitness values represent mean values per genotype. Mean flavone concentration per genotype was obtained through pooling of the genotype replicates before extraction of metabolites. Costs are identified in the absence of significant co-interactions (Flavones$_x$Treatment). Continuous lines refer to correlations with $P < 0.05$, and dotted lines refer to correlations with $P > 0.05$ from mixed effects models. *P* values on top of each plot refer to mixed effects models. Cya = cyanidin, Treat = treatment. N = 56.

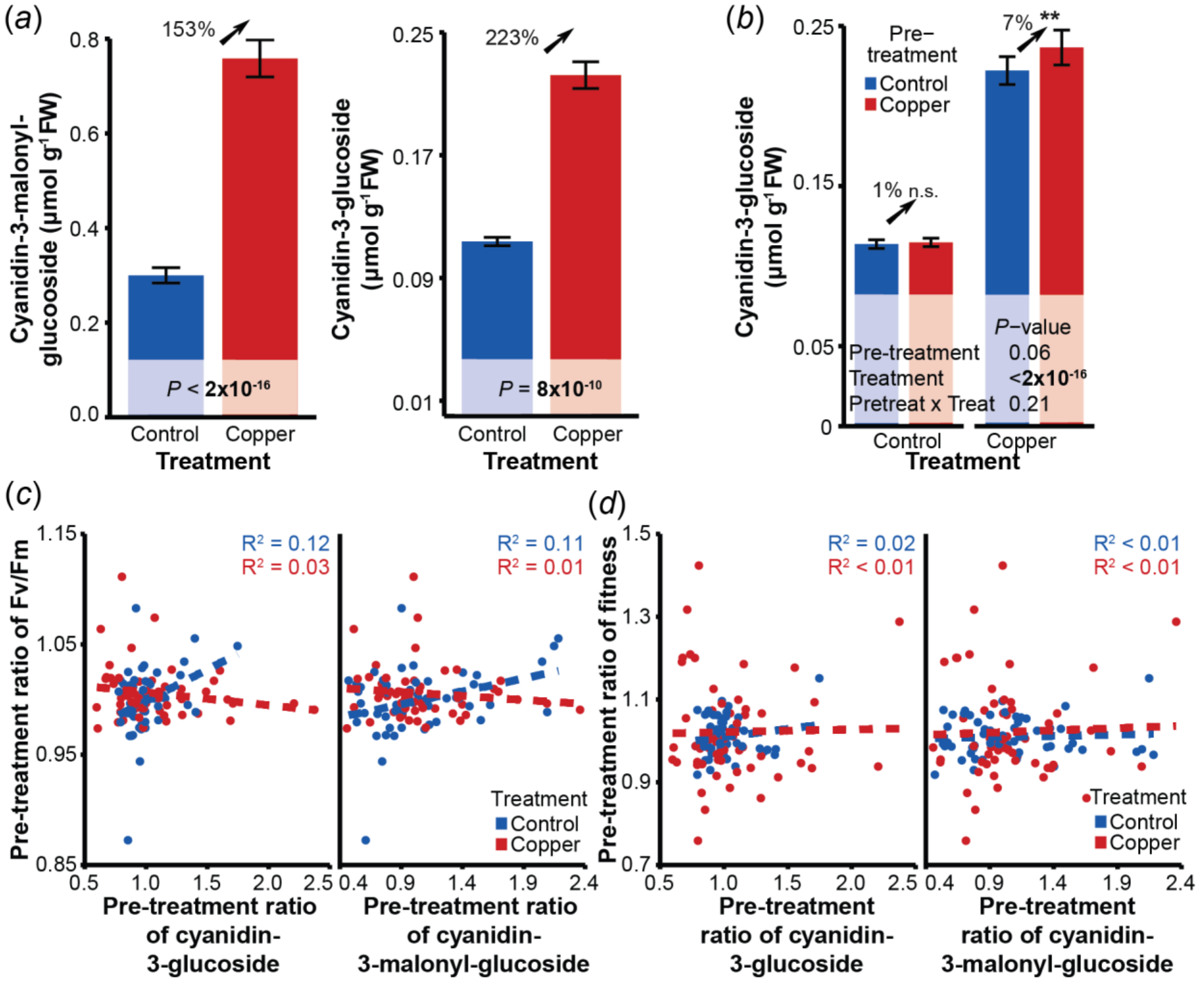


Figure S7. Copper excess induces high cyanidin concentrations that are transgenerationally retained elevated in the offspring, yet its transgenerational effects do not represent fitness nor phenotype advantages. (*a*) Copper excess enhanced cyanidin-3-malonyl-glucoside and cyanidin-3-glucoside concentrations. (*b*) Cyanidin-3-glucoside levels were transgenerationally retained elevated regardless of the presence or absence of recurrent copper excess. *P* values refer to mixed effects models. (*c*) The pre-treatment ratios of both cyanidins did not correlate with the pre-treatment ratios of photosystem II activity. *P* values refer to mixed effects models. For (*a*)-(*c*): Mean cyanidin concentrations per genotype were obtained through pooling of the genotype replicates before extraction of metabolites. (*d*) The pre-treatment ratios of cyanidins did not correlate with the pre-treatment ratios of plant fitness. Pre-treatment ratios of Fv/Fm and fitness refer to mean pre-treatment ratios per genotype. Dotted lines refer to correlations with *P* > 0.05 obtained from simple linear models. Pretreat = pre-treatment, Treat = treatment. N = 56.

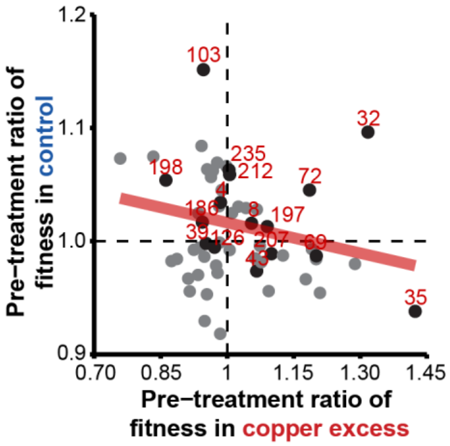


Figure S8. Genotypes in the repetition experiment represent the fitness trade-off effects. 16 genotypes (black dots with red letters) were selected to assess the reproducibility of transgenerational plasticity.

### Tables

Table S1. World-wide distributed genotypes of *Spirodela polyrhiza* used for the transgenerational experiments.

| Accession ID | Clone ID | Original Library | Population | Continent | Country | 16-genotype experiment |
|---|---|---|---|---|---|---|
| SP003 | 7379 | KlausApp | - | Asia | India | |
| SP004 | 7498 | KlausApp | America | North America | USA | Repeated |
| SP008 | 8683 | KlausApp | America | Africa | Kenya | Repeated |
| SP011 | 9242 | KlausApp | America | South America | Ecuador | |
| SP015 | 9510 | KlausApp | Europe | Africa | Mozambique | |
| SP026 | 9633 | KlausApp | Europe | Europe | Albania | |
| SP032 | 9505 | KlausApp | America | North America | Cuba | Repeated |
| SP035 | 0109 | KlausApp | SE Asia | Asia | China | Repeated |
| SP039 | 9650 | KlausApp | India | Asia | India | |
| SP043 | 9497 | KlausApp | India | Asia | India | Repeated |
| SP050 | 0090 | KlausApp | America | Asia | China | Repeated |
| SP055 | 9610 | KlausApp | Europe | Europe | Poland | |
| SP056 | 9256 | KlausApp | Europe | Europe | Finland | |
| SP059 | 9506 | KlausApp | India | Asia | India | |
| SP069 | 6731 | KlausApp | - | North America | USA | Repeated |
| SP072 | 5522 | KlausApp | SE Asia | Asia | China | Repeated |
| SP079 | 5513 | KlausApp | Europe | Europe | Germany | |
| SP087 | 9657 | Landolt | America | North America | Canada | |
| SP099 | 5524 | KlausApp | SE Asia | Asia | Thailand | |
| SP103 | | Simon&Martin | Europe | Europe | Switzerland | Repeated |
| SP110 | S21 | Simon&Martin | - | Europe | Switzerland | |
| SP111 | Ch1 | Sowjanya Sree | - | Asia | China | |
| SP115 | Kor3 | Sowjanya Sree | SE Asia | Asia | Korea | |
| SP117 | J2 | Sowjanya Sree | SE Asia | Asia | Japan | |
| SP126 | 8229 | Sowjanya Sree | SE Asia | Asia | Malaysia | Repeated |
| SP133 | 1059 | Sowjanya Sree | Europe | Asia | China | |
| SP141 | | YNQJ1 | SE Asia | Asia | China | |
| SP142 | | YNKM1 | SE Asia | Asia | China | |
| SP143 | | GZTR1 | SE Asia | Asia | China | |
| SP160 | | CSP17 | SE Asia | Asia | China | |
| SP165 | | CSP22 | SE Asia | Asia | China | |
| SP167 | | JXYT | SE Asia | Asia | China | |
| SP171 | | HNHY | SE Asia | Asia | China | |
| SP172 | | HNYZ | SE Asia | Asia | China | |
| SP178 | 0245 | ZH0245 | SE Asia | Asia | China | |
| SP179 | 0256 | ZH0256 | SE Asia | Asia | China | |
| SP180 | 0282 | ZH0282 | SE Asia | Asia | China | |
| SP181 | 0289 | ZH0289 | SE Asia | Asia | China | |
| SP183 | 0308 | ZH0308 | SE Asia | Asia | China | |
| SP185 | | SH5 | SE Asia | Asia | China | |

| | | | | | | |
|---|---|---|---|---|---|---|
| SP186 | | SH5 | SE Asia | Asia | China | Repeated |
| SP192 | Hgw | 17S | Europe | Europe | Germany | |
| SP194 | Wan | 18S | Europe | Europe | Germany | |
| SP197 | | Weihai5 | SE Asia | Asia | China | Repeated |
| SP198 | | | SE Asia | Asia | China | Repeated |
| SP207 | | Jiaonan5 | SE Asia | Asia | China | Repeated |
| SP208 | | | SE Asia | Asia | China | |
| SP210 | | | SE Asia | Asia | China | |
| SP212 | | Jining1 | SE Asia | Asia | China | Repeated |
| SP216 | | Hefeii | SE Asia | Asia | China | |
| SP221 | | UK2_01 | India | Asia | India | |
| SP223 | | UK2_07 | India | Asia | India | |
| SP224 | | UK2_10 | India | Asia | India | |
| SP225 | | NE1 | India | Asia | India | |
| SP227 | | NE3 | India | Asia | India | |
| SP235 | | JK5 | India | Asia | India | Repeated |

Accession ID: Code assigned in our group.

Clone ID: Code assigned in other publications or in the original library from other groups.

Population: Populations were identified from SNP analysis done by Wang Y, Duchen P, Chávez A, Sree KS, Appenroth KJ, Zhao H, Höfer M, Huber MXu S [1] and Xu S, Stapley J, Gablenz S, Boyer J, Appenroth KJ, Sree KS, Gershenzon J, Widmer AHuber M [2].

“-“ refers to unknown population as the sample was not sequenced.

Continent and Country: Geographical place were the samples were collected.

Table S2. Order of the 56 genotypes within the racks for the 56-genotype experiment.

| | Genotype | | | |
|---|---|---|---|---|
| Rack\Well | 1-2 | 3-4 | 5-6 | 7-8 |
| 1 | 11 | 15 | 26 | 32 |
| 2 | 50 | 69 | 87 | 32 |
| 3 | 171 | 224 | 181 | 207 |
| 4 | 172 | 8 | 87 | 225 |
| 5 | 185 | 186 | 194 | 223 |
| 6 | 117 | 126 | 133 | 115 |
| 7 | 178 | 11 | 99 | 227 |
| 8 | 142 | 143 | 160 | 141 |
| 9 | 50 | 26 | 110 | 180 |
| 10 | 35 | 39 | 43 | 50 |
| 11 | 197 | 198 | 210 | 212 |
| 12 | 126 | 180 | 143 | 171 |
| 13 | 35 | 56 | 223 | 115 |
| 14 | 186 | 210 | 8 | 224 |
| 15 | 192 | 208 | 3 | 11 |
| 16 | 99 | 103 | 110 | 111 |
| 17 | 59 | 194 | 160 | 111 |
| 18 | 103 | 15 | 179 | 235 |
| 19 | 43 | 186 | 212 | 126 |
| 20 | 179 | 180 | 181 | 178 |
| 21 | 111 | 133 | 160 | 172 |
| 22 | 165 | 167 | 171 | 172 |
| 23 | 43 | 192 | 198 | 208 |
| 24 | 55 | 35 | 99 | 72 |
| 25 | 59 | 110 | 183 | 142 |
| 26 | 216 | 221 | 224 | 225 |
| 27 | 72 | 165 | 3 | 216 |
| 28 | 227 | 235 | 192 | 208 |
| 29 | 223 | 216 | 227 | 197 |
| 30 | 198 | 179 | 185 | 167 |
| 31 | 72 | 79 | 87 | 183 |
| 32 | 32 | 55 | 143 | 183 |
| 33 | 69 | 141 | 197 | 133 |
| 34 | 55 | 56 | 59 | 69 |
| 35 | 212 | 181 | 194 | 225 |
| 36 | 185 | 39 | 117 | 210 |
| 37 | 4 | 79 | 167 | 221 |
| 38 | 115 | 141 | 165 | 178 |
| 39 | 3 | 4 | 8 | 207 |
| 40 | 142 | 79 | 103 | 117 |
| 41 | 39 | 15 | 56 | 4 |
| 42 | 26 | 207 | 235 | 221 |

Table S3. Order of the 16 genotypes within the racks for the 16-genotyoe experiment.

| | Genotype | | | |
|---|---|---|---|---|
| Rack\Well | 1-2 | 3-4 | 5-6 | 7-8 |
| 1 | 197 | 32 | 207 | 4 |
| 2 | 32 | 8 | 103 | 207 |
| 3 | 186 | 212 | 32 | 50 |
| 4 | 72 | 197 | 35 | 4 |
| 5 | 69 | 35 | 197 | 198 |
| 6 | 212 | 186 | 69 | 207 |
| 7 | 43 | 198 | 197 | 103 |
| 8 | 35 | 103 | 186 | 32 |
| 9 | 43 | 72 | 186 | 198 |
| 10 | 126 | 72 | 212 | 35 |
| 11 | 103 | 50 | 126 | 72 |
| 12 | 8 | 212 | 72 | 126 |
| 13 | 69 | 235 | 186 | 35 |
| 14 | 235 | 50 | 69 | 186 |
| 15 | 8 | 103 | 197 | 32 |
| 16 | 126 | 43 | 72 | 198 |
| 17 | 235 | 126 | 43 | 72 |
| 18 | 198 | 43 | 126 | 72 |
| 19 | 69 | 4 | 50 | 186 |
| 20 | 212 | 69 | 103 | 50 |
| 21 | 4 | 32 | 197 | 207 |
| 22 | 207 | 212 | 197 | 235 |
| 23 | 207 | 235 | 32 | 197 |
| 24 | 8 | 4 | 69 | 126 |
| 25 | 8 | 235 | 69 | 186 |
| 26 | 43 | 35 | 103 | 4 |
| 27 | 50 | 4 | 207 | 212 |
| 28 | 4 | 235 | 35 | 8 |
| 29 | 198 | 198 | 207 | 8 |
| 30 | 198 | 43 | 8 | 235 |
| 31 | 50 | 198 | 35 | 212 |
| 32 | 126 | 50 | 43 | 103 |

Table S4. Order in which the offspring of the 16-genotype experiment was placed to start the pre-treatment and treatment phases.

| | **Pre-treatment phase** | | **Treatment phase** | |
|---|---|---|---|---|
| Replicate | Copper | Control | Copper | Control |
| 1 | 1st offspring | 2nd offspring | 1st offspring | 2nd offspring |
| 2 | 1st offspring | 2nd offspring | 2nd offspring | 1st offspring |
| 3 | 1st offspring | 2nd offspring | 1st offspring | 2nd offspring |
| 4 | 1st offspring | 2nd offspring | 2nd offspring | 1st offspring |
| 5 | 2nd offspring | 1st offspring | 1st offspring | 2nd offspring |
| 6 | 2nd offspring | 1st offspring | 2nd offspring | 1st offspring |
| 7 | 2nd offspring | 1st offspring | 1st offspring | 2nd offspring |
| 8 | 2nd offspring | 1st offspring | 2nd offspring | 1st offspring |

Table S5. Genotypes that showed one replicate with smaller fitness than the 60-75% of the genotype median fitness under copper excess treatment.

| **Fitness lower than:** | **60%** | **65%** | **70%** | **75%** |
|---|---|---|---|---|
| ***P* (proportion test)** | 0.003 | 0.015 | 0.015 | 0.0025 |
| | SP15 | SP15 | SP15 | SP15 |
| | SP26 | SP26 | SP26 | SP26 |
| | SP32 | SP32 | SP32 | SP32 |
| | | SP72 | SP72 | SP72 |
| | | SP110 | SP110 | SP110 |
| | SP133 | SP133 | SP133 | SP133 |
| | SP143 | SP143 | SP143 | SP143 |
| **Pre-treatment:** | SP172 | SP172 | SP172 | SP172 |
| **Control** | | SP181 | SP181 | SP181 |
| | | | | SP185 |
| | SP194 | SP194 | SP194 | SP194 |
| | SP197 | SP197 | SP197 | SP197 |
| | | | | SP208 |
| | | | | SP221 |
| | SP223 | SP223 | SP223 | SP223 |
| | SP227 | SP227 | SP227 | SP227 |
| | | SP110 | SP110 | SP110 |
| **Pre-treatment: Copper** | | | SP142 | SP142 |
| | | SP179 | SP179 | SP179 |

### Formulas

Equation S1. Fitness as relative population growth rate [3] based on surface area of fronds.

$$\frac{\ln\text{ (final surface area)} - \ln\text{ (initial surface area)}}{\text{number of days}}$$

Equation S2. Pre-treatment ratios [4].

$$\frac{\text{plant fitness or phenotype of copper pre} - \text{treated plants}}{\text{mean (plant fitness or phenotype of control pre} - \text{treated plants)}}$$

Equation S3. Resistance ratios.

$$\frac{\text{plant fitness under first time stress}}{\text{mean (plant fitness under control conditions)}}$$

### Methods

Methods S1. Sterilization of plants

To surface sterilize the plants, we first treated each *S. polyrhiza* genotype with diluted DanKlorix (sodium hypochlorite containing disinfection solution, CP GABA GmbH, Hamburg, Germany) and subsequently with N-medium containing 100 µg $mL^{-1}$ cefotaxime sodium salt (Duchefa Biochemie, Haarlem, Netherlands) and 50 mM sucrose.

To prepare the media, we added 17.1 g of sucrose to 2L of N-medium and autoclaved the media. We then added 1µl cefotaxime stock (100 mg $mL^{-1}$) per 1 mL medium under the sterile bench. We dissolve the sodium hypochlorite mix in distilled water in a proportion of 1:10 (5 mL DanKlorix + 45 mL $dH_2O$).

To sterilize the plants under the sterile bench, we placed 3-5 fronds within a falcon tube containing the dissolved sodium hypochlorite and incubated them for 10-15 min with occasional inversion of the tubes. Afterwards, we transferred the fronds to sterilized 100 mL flasks containing 50 mL of N-medium with cefotaxime and sucrose and let the plants regenerate at 26°C, 120 µmol photons $m^2$ $s^{-1}$ for minimum eight days. Regenerated plants were transferred into 250 mL flask with 150-180 mL N-medium for pre-cultivation. Recovered plants were placed into sucrose N-medium before experiments to corroborate their sterility.

Methods S2. Parameter within the IMAGING-PAM MAXI version

To measure the photosystem II efficiency, we used the software of IMAGING-PAM M-Series MAXI version, ImagingWinGigE V2.47+, to set all the parameters. The intensity of the saturating pulse to obtain the $F_v/F_m$ ratios was of 3500 µmol $m^{-2}$ $s^{-1}$, for a width of 720 ms. The setting was:

Set gain == 5
Set damping == 2
Set measuring light == 3
Set measuring frequency == 4
Set saturation light == 10

Once the script was executed, we waited 10 seconds in before and after measuring $F_v/F_m$ ratios.

Methods S3. Flavonoid analysis

To extract the metabolites, we used 30 mg of flash-frozen and grinded plant material. For the extraction, we applied into the samples acidified methanol (MeOH:water:formic acid 15:4:1 v/v/v). The whole extraction followed similar steps as described in Malacrinò A, Böttner L, Nouere S, Huber M, Schäfer MXu S [5], with focus on the flavonoids.

In details, the flavones were analysed in a 1:100 dilution of the extract in an aqueous mixture of stable isotope labelled amino acid (Algal amino acid mixture-$^{13}C$,$^{15}N$, Sigma) by LC-MS. Chromatographic separation was done on a Shimadzu Nexera X3 LC-System. The system was equipped with an Agilent 1290 infinity II inline filter (0.3 µm) and a ZORBAX RRHD Eclipse XDB-C18 column (3x50 mm, 1.8 µm; Agilent Technologies). The mobile phase comprised 0.05% formic acid (Fisher Chemical), 0.1% acetonitrile (Fisher Chemical) in water as Solvent A and methanol (Fisher Chemical) as Solvent B in gradient mode with the elution profile: 0–1.5 min, 2% B; 1.5–3.5 min, 2%–100% B; 3.5–4.5 min, 100% B; 4.5–5 min, 100%-2% B and 5-6 min, 2 % B, with a flow rate of 500 µL/min. The column oven was set to 42 °C. The measurements were performed on a Shimadzu LCMS-8060 equipped with an ESI source, which was operated in multi-reaction-monitoring (MRM) modus. Settings were as follows: Nebulizing Gas Flow: 3 L/min; Heating Gas Flow: 10 L/min; Drying Gas Flow: 10 L/min; Interface Temperature: 300 °C; DL Temperature 250 °C; Heat Block Temperature: 400 °C; CID Gas: 270 kPa; Q1 Resolution: Unit; Q3 Resolution: Unit.

The detector settings (MRM-settings and retention times) were:

| Analyte | RT [min] | Q1 [m/z] | Q3 [m/z] | Dwell time [ms] | CE [V] | Q1/Q3 Pre Bias [V] | ISTD |
|---|---|---|---|---|---|---|---|
| Apigenin-7-O-glucoside | 3.83 | (+) 431,10 | 268,10 | 32 | -35 | -21 / -17 | $^{13}C_9$, $^{15}N_1$-Phe (2.25) |
| | | (+) 431,10 | 269,20 | 32 | -28 | -21 / -17 | |
| Apigenin-8-C-glucoside | 3.69 | (+) 431,10 | 311,20 | 32 | -24 | -30 / -21 | $^{13}C_9$, $^{15}N_1$-Phe (2.37) |
| | | (+) 431,10 | 283,25 | 32 | -35 | -30 / -19 | |
| Luteolin-7-O-glucoside | 3.73 | (+) 447,09 | 285,20 | 32 | -26 | -22 / -19 | $^{13}C_9$, $^{15}N_1$-Phe (1.62) |
| | | (+) 447,09 | 284,20 | 32 | -37 | -22 / -18 | |
| Luteolin-8-C glucoside | 3.63 | (+) 447,09 | 327,15 | 32 | -24 | -30 / -22 | $^{13}C_9$, $^{15}N_1$-Phe (1.26) |
| | | (+) 447,09 | 357,15 | 32 | -21 | -30 / -24 | |
| $^{13}C_9$, $^{15}N_1$-Phe | 2.52 | (+) 176.11 | 129.25 | 247 | -15 | -20/-20 | |

RT: retention time.

CE: collision energy.

ISTD: internal standard incl. response factor (nAnalyte = x * nISTD) in brackets.

Qualifiers are highlighted in grey.

On the other hand, anthocyanins were analysed by HPLC-PDA, thereto the chromatographic separation was done on a Shimadzu Nexera XR LC-System equipped with an EC 4/3 Nucleodur Sphinx RP pre-column (5 µm, Macherey-Nagel) and a Nucleodur Sphinx RP column (250x4.6 mm, 5µm, Macherey-Nagel). The mobile phase comprised 0.2% formic acid (Fisher Chemical), 0.1% acetonitrile (Fisher Chemical) in water as Solvent A and acetonitrile (Fisher Chemical) as Solvent B in gradient mode with the elution profile: 0–8 min, 10-21% B; 8-18 min, 21%–49% B; 18–18.1 min, 49-100% B; 18.1–19.0 min, 100% B; 19.0–19.1 min, 100%-10% B and 19.1-24 min, 10 % B, with a flow rate of 1300 µL/min.. The column oven was set to 20 °C. Measurement was performed with a PDA detector.

To quantify the anthocyanins, we used an external dilution curve of the molar quantity of cyanidin-3-O-glucoside. Both cyanidins had the following retention time and absorption wavelength:

| Analyte | Retention time [min] | Absortion wave lenght [nm] |
|---|---|---|
| Cyanidin-3-O-glucoside | 7.055 | 517 |
| Cyanidin-3-O-(6-O-malonyl-beta-glucoside) | 9.41 | 517 |
